# AMPA receptor activation within the prelimbic cortex is necessary for incubated cocaine-craving

**DOI:** 10.1101/2025.05.21.655393

**Authors:** Laura L. Huerta Sanchez, Mirette G. Tadros, Hoa H.T. Doan, Sylvie V. Vo, Sanil R. Chaudhari, Taylor L. Li, Peter B. James, Audrey Y. Na, Fernando J. Cano, Tod E. Kippin, Karen Szumlinski

## Abstract

The incubation of craving is a behavioral phenomenon in which cue-elicited craving increases during a period of drug abstinence. Incubated cocaine-craving is associated with increased extracellular glutamate within the medial prefrontal cortex (mPFC) and this release, particularly within the prelimbic (PL) subregion, is necessary for incubated cocaine-craving. A potential candidate mediating these incubation-driving effects of glutamate release within the PL are alpha-amino-3-hydroxy-5-methyl-4-isoxazolepropionic acid receptors (AMPARs). To investigate the role of mPFC AMPARs in incubated craving, male and female Sprague-Dawley rats were trained to self-administer cocaine for 6 h/day for 10 consecutive days. Either during early or later withdrawal, rats were infused intra-PL with the AMPAR antagonist NBQX (0 or 1 µg/0.5 µl per side), followed by 30-min tests for cue-reinforced responding. Immunoblotting was also conducted to relate the expression of incubated cocaine- and sucrose-craving to AMPAR subunit expression within mPFC subregions. Intra-PL NBQX blocked incubated craving expressed in late, but not early, withdrawal. No incubation-related changes in AMPAR subunit expression were detected within the PL or IL of rats of either sex and no estrus-associated changes in subunit expression were detected in female rats exhibiting incubated cocaine-craving. In contrast, elevated GluA1 expression was observed within the IL of male rats exhibiting an incubation of sucrose-craving. Together, these findings indicate a necessary role for AMPARs within the PL in driving incubated cocaine-craving and suggest that AMPARs located within the IL may be involved also in sucrose-craving selectively in males.

## Introduction

Cocaine use disorder (CUD) is a chronic, relapsing disorder that leads to devasting behavioral and physical health complications. In 2023, approximately 5 million people in the United States, aged 12 or older, reported using cocaine in the past 12 months. Further, among these individuals, 1.3 million people were diagnosed with a cocaine use disorder in the past year (SAMHSA 2023). CUD is characterized by a high occurrence of relapse, especially during protracted withdrawal. One factor driving relapse is re-exposure to drug-associated cues. Drug cues induce cravings that can intensify or “incubate” over a period of abstinence, rendering those in recovery more sensitive to the motivational pull of drug-associated cues to drive relapse (Grimm et al., 2001). This so-called “incubation of craving” phenomenon has been demonstrated in both humans and in laboratory animals, the latter of which enables direct investigation of the neurobiology underlying incubated drug-craving (c.f., Chow et al., 2025). As summarized in a recent review (Chow et al., 2025), despite nearly two decades of research focused on the neurobiology of incubated drug-craving, the precise neuroanatomy and molecular mechanisms underpinning incubated drug-craving remain to be elucidated. While a large body of animal research points to the nucleus accumbens (NAc) as an important neural locus in the circuitry underpinning drug craving and it’s incubation during protracted drug abstinence (Chow et al., 2025; Dong et al., 2017), craving induced by exposure to drug-related cues reliably increases prefrontal cortex (PFC) activity in humans with substance use disorders (e.g., Devoto et al., 2020; Goldstein & Volkow, 2011; Mohd Nawawi et al., 2024) and increases indices of cellular activity within the medial aspect of the PFC (mPFC) in animal models of incubated drug-craving (e.g., Huerta Sanchez et al., 2023; Koya et al., 2009; Miller et al., 2016; Szumlinski et al., 2018).

The mPFC is composed of the prelimbic (PL) and infralimbic cortices (IL) and both mPFC subregions project to, and receive excitatory glutamate projections from, many reward-related regions, including the NAc and amygdala (Kalivas et al., 2005; Manoocheri and Carter, 2022; Sesack et al., 1989). Supporting a link between incubated cocaine-craving and glutamate hyperactivity within the mPFC, rats expressing incubated cocaine-seeking exhibit a cue-elicited rise in extracellular glutamate within the mPFC that is not apparent in rats tested in early withdrawal (Shin et al., 2016). Directly implicating this glutamate release as a key driver of incubated cocaine-craving, infusion of a group 2/3 metabotropic glutamate autoreceptor agonist into the PL completely blocks incubated cue-elicited cocaine-seeking, with a measurable, but less robust, effect observed also when the agonist was infused into the IL (Shin et al., 2018). Our neuropharmacological findings contrast with those obtained under the extinction-reinstatement model of cocaine-seeking in which a dorsal-ventral dichotomy appears to exist with respect to how these two mPFC subregions modulate cocaine-seeking behavior (Kalivas, 2009; Kalivas and McFarland, 2003; LaLumiere et al., 2012; Peters et al., 2008). However, our observation that glutamate release within both PL and IL subregions contribute to driving incubated cocaine-seeking (Shin et al., 2018) aligns with optogenetics data implicating the unsilencing of synapses within both PL-NAc and IL-NAc projections in the development of incubated cocaine-craving (Ma et al., 2014).

Having established that glutamate release within the mPFC is required for the expression of incubated cocaine-craving (Shin et al., 2018), the question arises as to which postsynaptic glutamate receptors within the mPFC might be mediating the “incubation-driving” effects of cue-elicited glutamate release? Potential receptor candidates may be one or more of the ionotropic glutamate receptors (iGluRs) that rapidly depolarize neurons upon stimulation and mediate “fast” synaptic transmission (c.f., Diering and Huganir, 2018). Of the three iGluRs, alpha-amino-3-hydroxy-5-methyl-4-isoxazolepropionic acid receptors (AMPARs) have received considerable attention in the context of incubated drug-seeking, particularly those located within the NAc (c.f., Chow et al., 2025; Loweth et al., 2014). AMPARs are tetrameric ion channels, comprised of GluA1, GluA2, GluA3 and GluA4 subunits. In forebrain, most AMPARs consist of GluA1 and GluA2 subunits, gate Na^+^ influx and are impermeable to Ca^2+^(CI-AMPARs) (Boudreau et al., 2007; Kourrich et al., 2007; Conrad et al., 2008; Reimers et al., 2011). Pharmacological inhibition of AMPARs using the general receptor antagonists CNQX or NBQX directly into the NAc reduces cocaine-seeking during protracted withdrawal (e.g., Di Ciano et al., 2001; Doyle et al., 2014; Lynch et al., 2021). However, during protracted withdrawal from daily extended-access to intravenous cocaine, the cell surface expression of the GluA1 subunit is increased within the NAc, along with the synaptic insertion of Ca^2+^-permeable AMPARs (CP-AMPARs) that lack the GluA2 subunit. Supporting a functional role for CP-AMPAR insertion in incubated cocaine-craving, an intra-NAc infusion of the CP-AMPAR-selective antagonist Naspm blocks cocaine-craving, but only in rats tested in protracted withdrawal when CP-AMPAR expression is high (Conrad et al., 2008; Loweth et al., 2014). The Naspm effect is sex-independent as it is apparent in both male (Loweth et al., 2014) and female rats (Kawa et al., 2022). Aligning with these data, optogenetics studies implicate the insertion of CP-AMPARs in the maturation of silent synapses within IL-NAc projections during protracted cocaine withdrawal, while the insertion of CI-AMPARs contribute to the maturation of silent synapses within PL-NAc projections purported to drive incubated cocaine-seeking behavior (Ma et al., 2014). While a similar insertion of CP-AMPARs are reported to occur within the PFC of mice injected repeatedly with cocaine (Ruan and Yao, 2021), to the best of our knowledge, the functional relevance of AMPARs within mPFC subregions for the expression of incubated drug-craving following a period of voluntary self-administration has not been investigated directly.

As a first-pass examination of the role for mPFC AMPARs in incubated cocaine-craving, the current studies examined the effects of an intra-PL infusion of the AMPAR antagonist NBQX on cue-elicited craving expressed by female and male rats during early versus later withdrawal from a history of long-access (6 h/day) intravenous cocaine self-administration. Immunoblotting for GluA1 and GluA2 subunits was also conducted on tissue from mPFC subregions to compare the relationship between AMPAR subunit levels and the expression of incubated cocaine-versus sucrose-craving and to determine how estrous phase might influence AMPAR subunit expression during incubated craving in female rats. To the best of our knowledge, this study is the first to confirm that AMPAR activation within the PL subregion is required for the expression of incubated cocaine-craving in both female and male rats. Further, we show that the profile of GluA1 and GluA2 expression within whole-cell homogenates from mPFC subregions is distinct between rats expressing incubated cocaine-versus sucrose-seeking of relevance to our neurobiological understanding of these behavioral phenomena.

## Materials and Methods

### Subjects

Adult male (250-275g) and female (225-250g) Sprague Dawley rats (Charles River Laboratories, Hollister, CA) were housed in a colony room under 12-h reverse light cycle conditions (lights off: 10:00 am). Following arrival, rats were allowed to acclimate to the colony room for 48 h and were given *ad libitum* access to food and water throughout the study. All procedures were approved by the Institutional Animal Care and Use Committee of the University of California, Santa Barbara under protocol number 829 and were consistent with the guidelines of the NIH *Guide for Care and Use of Laboratory Animals*. Note that the rats employed in the immunoblotting study of incubated sucrose-craving were the same rats as those employed in Cano et al. (2025). As such, no new animals were required to conduct this experiment.

### Surgery

Under isoflurane anesthesia (4% induction, 1-3% maintenance; Covetrus, Portland, ME), rats were implanted with bilateral guide cannulae (P1 Technologies, Roanoke, VA) aimed above the PL subregion of the mPFC (AP: +3.0; ML: ±0.75, DV: -2.00 mm from Bregma) and secured to the skull with four stainless steel screws (Specialty Tool, Goleta, CA) and dental acrylic. Rats that were slated to undergo cocaine self-administration were also implanted with a chronic polyurethane catheter (12 cm long; 0.023 inner diameter, 0.038 in outer diameter; Instech Laboratories, Plymouth Meeting, PA) into the right jugular vein and ran subcutaneously over the shoulder to a back incision. The catheter was then secured to a 22-gauge guide cannula (P1 Technologies, Roanoke, VA) in a rat infusion harness (Instech Laboratories, Plymouth Meeting, PA) and capped to protect against infection. Following this procedure, catheters were flushed with 0.1 ml of sterile cefazolin (100 mg/ml) and 0.1 ml of sterile heparin (70 U/ml). Rats slated to be tested for cocaine-craving on WD1 underwent both surgeries on the same day, 7 days prior to the first cocaine self-administration session. Rats slated to be tested on WD30 underwent the IV catheter implantation surgery 5 days prior to cocaine self-administration training procedures and then underwent the intracranial implantation surgery 7 days prior to their test on WD30. Rats were monitored postoperatively for 4-7 days under which rats received subcutaneous Meloxicam (2 mg/kg) once a day for the first 2 postoperative days for pain and daily injections of cefazolin and heparin to maintain catheter patency. To ensure catheter patency prior to cocaine self-administration training, rats were injected IV with 0.1 ml of sodium Brevital (10 mg/ml).

### Cocaine and Sucrose Self-Administration Procedures

CP-AMPAR accumulation and increased glutamate transmission within the NAc appear to require long-access cocaine self-administration procedures to manifest (Purgianto et al., 2013). As our prior immunoblotting report failed to detect changes in AMPAR subunit expression following short-access cocaine self-administration procedures (Huerta Sanchez et al., 2023), the rats in the present study were trained to self-administer intravenous cocaine (0.25 mg per 0.1 ml saline infusion; MilliporeSigma, Burlington, MA) over 10 once-daily 6-h sessions under an FR1 schedule of reinforcement with a 20-sec time-point. As detailed in Cano et al. (2025), rats in the sucrose-craving study were trained to respond for delivery of a 45 mg banana-flavored sucrose pellet (BioServ, Flemington, NJ) under comparable conditions. In both cases, each press of the “active” lever resulted in a 20-second tone and light stimulus complex (78 dB, 2kHz) signaling the reinforcer delivery. Rats were not able to receive additional infusions or pellets during the cue presentation. To provide a baseline for protein expression, a group of cocaine-naive controls were included in the cocaine immunoblotting study (Controls) that only received the 20-second tone-light stimulus when they depressed the active lever (i.e., no primary reinforcer was available). In all cases, depression of the “inactive” lever produced no programmed consequences. On the first day of IV cocaine self-administration, rats were capped at 100 infusions to prevent overdose. While the rats slated for the immunoblotting studies did not undergo any lever-press training prior to the start of cocaine self-administration procedures, the rats slated for the neuropharmacological studies were first trained to lever-press for the sucrose pellets during two 6-h sessions prior to surgery to engender more reliable subsequent cocaine self-administration behavior, as conducted in previous microinjection studies by our group (e.g., Ben-Shahar et al., 2013; Szumlinski et al., 2019).

### Test for Cue-Elicited Cocaine- and Sucrose-Craving

At early or later withdrawal time-points, (WD1 or WD3 and WD30-31, respectively) rats were placed back into their assigned operant chambers to undergo tests for cue-elicited cocaine- or sucrose-craving. During this test, an active lever press resulted in the presentation of the same 20-second tone and light stimulus complex as experienced during self-administration training, but no cocaine or sucrose delivery. There were no programmed consequences following the depression of the inactive lever. For the rats slated for the immunoblotting studies, this test was 2-h long and tissue was extracted immediately following the end of the test session (see below). For the rats slated for the neuropharmacological study, the test was 30-min long and conducted immediately following the microinjection (see . Incubated cocaine- and sucrose-craving were defined as a statistically significant increase in active lever presses emitted by rats tested in later withdrawal (e.g., WD30 or WD31) versus early withdrawal (WD1 or WD3, depending on the study).

### Microinjection

To examine for the role of AMPARs within the PL in incubated cocaine-craving, the non-subunit selective AMPA antagonist 2,3-Dioxo-6-nitro-1,2,3,4-tetrahydrobenzo[*f*]quinoxaline-7-sulfonamide (NBQX; Tocris, Minneapolis, MN) was dissolved in 1% DMSO and infused at a dose of 1 µg/side, which is comparable to that employed in other studies of drug-induced behavior (Russell et al 2016; Biondo et al., 2005) and 1% DMSO served as the control infusion (VEH). On WD1 or WD30, rats were microinjected bilaterally at a rate of 0.5 µl/minute for 1 minute with either NBQX or VEH (total infusion volume/side = 0.5 µl) and microinjectors were left in place for an additional 30 sec prior to removal. Immediately following the microinjection, rats underwent the 30-minute cue test described above. To assess for any potential effects of NBQX on the consolidation of learning, rats were tested again the following day with no further microinjection (Cue Test 2; WD31). Once both tests were completed, rats were sacrificed, brains were extracted, fixed in 4% paraformaldehyde and sectioned (30 μm thick) for histological verification of microinjector placement using Nissl staining procedures.

### Immunoblotting

Following the 2-h cue-elicited cocaine-seeking test conducted on WD3 or WD30, brains were extracted and the PL and IL subregions of the mPFC were dissected over ice for immunoblotting. As we wanted the present results to be as comparable to prior immunoblotting studies of incubated cocaine-craving, the methods used for whole-cell tissue homogenate preparation, detection and quantification followed similar procedures as those used previously (Chiu et al, 2021; Huerta Sanchez et al., 2023). Due to the large number of experimental groups included in the cocaine immunoblotting study, the tissue was processed separately for male and female rats. As fewer groups were tested in the sucrose immunoblotting study (Cano et al., 2025), tissue from males and females were run concurrently on the same gels. To quantify AMPAR subunit expression within our samples, anti-rabbit GluA1 (1:500; Millipore; AB1504) and anti-mouse GluA2 (1:1000 dilution; Synaptic Systems; 182 111) primary antibodies were used. Calnexin expression was used to control for protein loading and transfer (anti-rabbit Calnexin primary antibody 1:1,000 dilution; Enzo Life Sciences; ADI-SPA-860). Following primary incubation, membranes were washed with TBST and incubated in either a goat anti-rabbit IRDye 800 CW secondary antibody (1:10,000 dilution; Li-Cor; 925-3221) or a goat anti-mouse IRDye 680RD secondary antibody (1:10,000 dilution; Li-Cor; 925-68070). Membranes were then washed for a second time and imaged in an Odyssey Fc Infrared Imaging System (Li-Cor Biosciences, Lincoln, NE, USA). Protein expression was quantified using Image Studio. Raw values for each band were normalized to their corresponding calnexin signal and then to the average value of the control group (i.e Control-WD3 for the cocaine study and WD1-males for the sucrose study). Blots exhibiting anomalies were excluded from the final statistical analysis of the data.

### Vaginal Cytology

Evidence suggests that the magnitude of incubated cocaine-craving varies as a function of the estrus cycle (Corbett et al., 2021; Kerstetter et al., 2008; Nicholas et al. 2019) and in hippocampus, the synaptic insertion of CP-AMPARs during inhibitory avoidance learning varies as a function of estrous cycle (Tada et al., 2015). Thus, we monitored each female’s estrous cycle via vaginal swabbing following each cue test. Vaginal samples were collected by gently swabbing the vaginal canal with a cotton-tipped applicator, soaked in sterile saline. Samples were then transferred onto glass microscope slides, sprayed with a fixative, and stained with Giemsa using standard procedures. The estrous stage was determined based on the presence and morphology of cells. Each sample was categorized into one of four estrous phases: proestrus, estrus, metestrus and diestrus. Proestrus can be recognized by the abundant presence of small nucleated epithelial cells, and estrus by the abundant presence of non-nucleated cornified epithelial cells. Metestrus is identified by the presence of approximately equal amounts of both small and big nucleated epithelial cells, non-nucleated cornified epithelial cells, and neutrophils. Diestrus is characterized by a low cell density and the presence of neutrophils, with occasional nucleated and almost no cornified epithelial cells. Given the relatively low number of female rats in metestrus, the data from the rats in metestrus and diestrus were combined for data analysis as conducted in prior studies (Nicolas et al., 2019).

### Statistical Analysis

Data was analyzed using analyses of variance (ANOVAs) to examine for behavioral and biochemical outcomes associated with incubation. For the study of the effects of NBQX on incubated cocaine-craving, the average number of active and inactive lever presses emitted during the cue tests were analyzed using a Treatment (WD1-VEH, WD30-VEH, WD30-NBQX) X Sex ANOVA. The data from the study of the effects of NBQX on responding in early withdrawal were analyzed using a Treatment (VEH vs. NBQX) X Sex ANOVA. For the immunoblotting study of incubated cocaine-craving, the data were expressed as a percentage of the average of the two or three Control-WD3 animals on each gel and analyzed using a Group (Control vs. Cocaine) X Withdrawal ANOVA, separately for male and female rats. The behavioral data from this immunoblotting study were also analyzed using Group X Withdrawal ANOVAs, separately for males and females for consistency. As the immunoblotting study of incubated sucrose-craving did not include a sucrose-naive control, the samples from males and females could be immunoblotting concurrently on the same gel. As such, these data were expressed as a percentage of the average of the three Male WD1 rats on each gel and analyzed using a Sex X Withdrawal ANOVA, as conducted previously (Cano et al., 2025). For the examination of the effect of estrous phase on behavior and protein expression within female cocaine-experienced rats, the data were analyzed using a Phase (estrus, diestrus, proestrus; metestrus females combined with diestrus) X Withdrawal ANOVA. Significant main effects or interactions in all analyses were further investigated with t-tests or tests for simple effects. Outliers were identified and excluded from the analyses using the ± 1 × IQR rule, however, in instances where too many outliers were identified, we adopted the ± 3 × IQR rule to ensure that only the most extreme outliers were removed. Alpha was set to 0.05 for all analyses with the exception of the analyses of estrous phase influence on behavior and protein expression in which alpha was set to 0.1 as we had *a priori* predictions that: (1) AMPAR subunit expression would vary with estrous cycle phase (Tada et al., 2015) and (2) incubated cocaine-craving would be highest in female rats in estrus (e.g., Kerstetter et al., 2008). IBM SPSS Statistics software (version 29.0 for Macintosh) was used for all statistical tests, and GraphPad Prism software (version 9.3.1 for Macintosh) was used to create all graphs.

## Results

### Intra-PL NBQX lowers incubated cocaine-craving

The results pertaining to the average behavior of the rats over the course of the last 3 days of the cocaine self-administration phase of the study are presented in **Table 1 (Expt. 1)**. Analyses of the number of active lever-presses [F(5,33)=0.014, p=0.906], inactive lever-presses [F(5,33)=2.457, p=0.127] and reinforcers earned [F(5,33)=0.079, p=0.861] did not indicate any significant group differences at the outset of testing. Further, on neither cue test day were sex differences in responding apparent on the active [F(5,29)<2.419, p’s>0.459] or inactive lever [F(5,30)<2.712, p’s>0.111]. As such the data were collapsed across the sexes for visualization of the NBQX effect on incubated responding (**Figure 1**).

**Figure 1:**
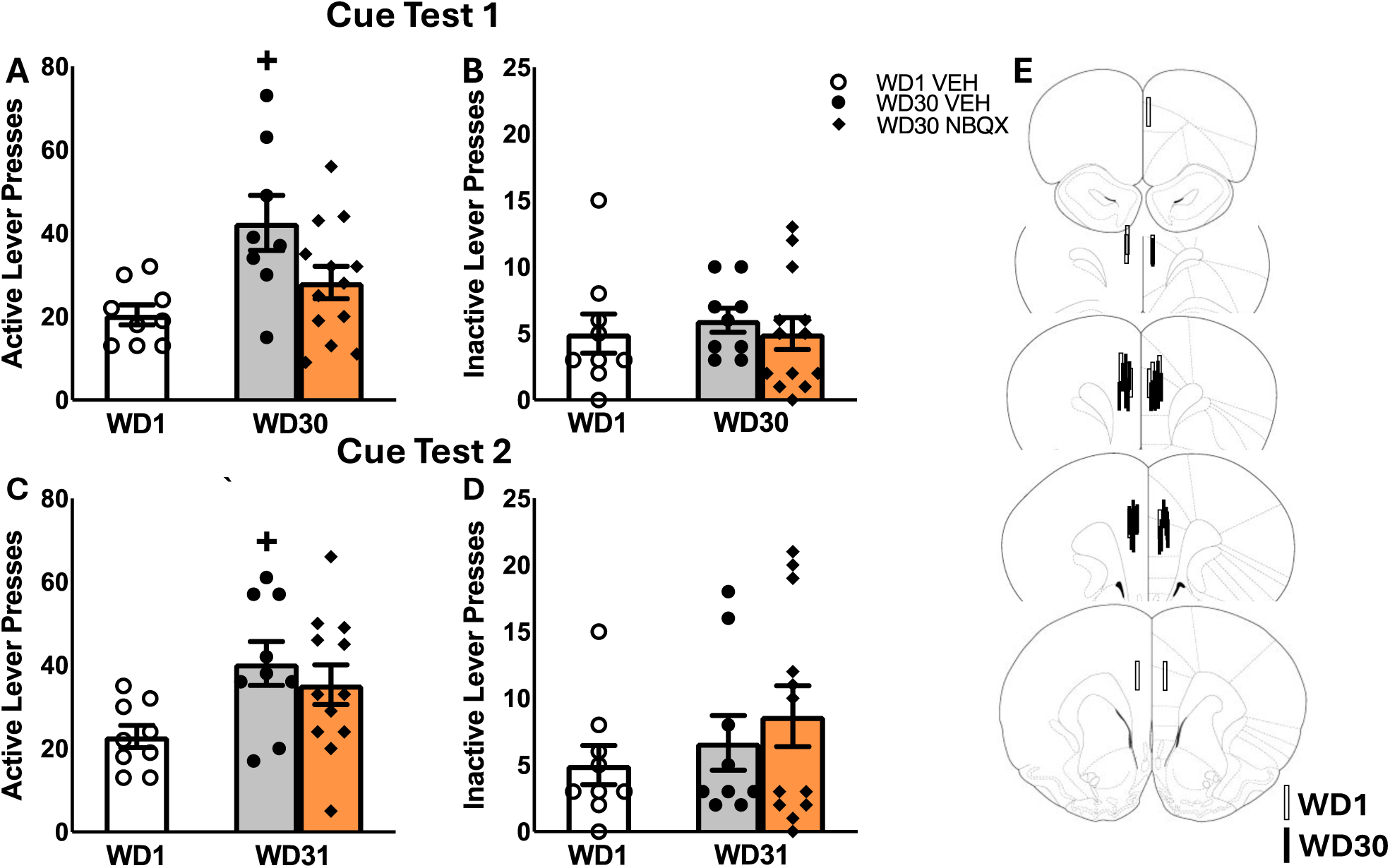
Summary of the effects of an intra-PL infusion of NBQX (1 *μ*g/side) or vehicle (VEH) on active and inactive lever-responding during tests for incubated cocaine-craving conducted either immediately following microinfusion (A,B) and during a second test conducted 24 h later (C,D). Note: As no sex differences in responding were detected, the data is collapsed across male and female rats for better visualization of the NBQX effect. The data represent the means ± SEMs of the number of individual rats indicated. (E) Cartoon depicting the placements of the microinjectors within the PL. +p<0.05 vs. WD1 (withdrawal day 1).

**Table 1:** Summary of the data from the self-administration training phases of each experiment described in this report. The data represent the means ± SEMs. Note that the data from Expt. 4 are derived from Cano et al. (2025).

|  | Active | Reinforcer | Inactive |
| --- | --- | --- | --- |
| **Exp. 1 – NBQX effect on incubated cocaine-craving** |  |  |  |
| WD1 VEH | 230.3±148.09 | 107.27±24.14 | 7.47±1.49 |
| WD30 VEH | 133.03±45.63 | 79.5±15.86 | 8.47±2.16 |
| WD30 NBQX | 159.67±69.58 | 88.28±13.22 | 22.49±9.64 |
| **Exp. 2 – NBQX effect in early cocaine withdrawal** |  |  |  |
| WD1 VEH | 100.26±17.41 | 77.86±14.58 | 13.85±5.56 |
| WD1 NBQX | 99.66±11.41 | 89.03±9.89 | 2.73±1.06 |
| **Exp. 3 – Immunoblotting: Incubated Cocaine-Craving** |  |  |  |
| **Male:** |  |  |  |
| WD3 Control | 26.55±8.72 | 18.83±1.99 | 12.87±5.54 |
| WD3 Cocaine | 64.06±11.88 | 52.53±2.42 | 6.23±9.48 |
| WD30 Control | 18.15±5.21 | 14.92±1.65 | 10.35±3.90 |
| WD30 Cocaine | 81.4±12.47 | 74.47±2.35 | 7.48±12.37 |
| **Female:** |  |  |  |
| WD3 Control | 28.17±11.07 | 7.9±1.91 | 5.47±1.14 |
| WD3 Cocaine | 70.4±9.69 | 64.93±0.73 | 3.73±9.69 |
| WD30 Control | 7.73±1.30 | 13.85±0.59 | 2.73±6.96 |
| WD30 Cocaine | 70.74±7.67 | 65.15±11.18 | 18.15±5.71 |
| **Exp.4 – Immunoblotting: Incubated Sucrose-Craving** |  |  |  |
| **Male:** |  |  |  |
| WD1 | 197.91±11.40 | 170.64±11.54 | 12.33±2.68 |
| WD30 | 170.57±10.08 | 140.51±7.59 | 15.55±3.02 |
| **Female:** |  |  |  |
| WD1 | 184.4±12.00 | 160.22±9.04 | 9.95±3.31 |
| WD30 | 176.55±10.43 | 140.52±9.26 | 16.64±4.52 |

A comparison of the number of active (**Figure 1A**) and inactive (**Figure 1B**) lever-presses emitting during a 30-min test for cue-reinforced responding indicated a significant Treatment effect for active lever responding [for active lever: F(5,30)=5.047, p=0.013; for inactive lever: F(5,33)=0.151, p=0.860]. As expected, VEH-infused rats tested on WD30 emitted more active lever-presses than the WD1 controls [t(16)=3.158, p=0.006], indicative of incubated cocaine-craving in the WD30 controls. In contrast, active lever-responding did not differ between NBQX-infused rats tested on WD30 versus VEH-infused rats tested in either early [t(21)=1.264, p=0.220] or later withdrawal [t(19)=.435, p=0.060], indicating that intra-PL NBQX was sufficient to lower cue-reinforced responding on WD30 to block the expression of incubated cocaine-craving.

When tested the next day in the absence of any further pretreatment, we again detected a significant Treatment effect for the number of active lever-presses (**Figure 1C,D**) [for active lever: F(5,29)=4.918, p=0.015; for inactive lever: F(5,30)=.837, p=0.444]. Consistent with our prior studies (e.g., Ben-Shahar et al., 2013; Chiu et al., 2021; Szumlinski et al., 2019), incubated cocaine-craving persisted in VEH-infused rats tested on WD31 [t(16)=3.464, p=0.003]. Cue-reinforced responding of NBQX-pretreated rats did not differ from their VEH-infused counterparts tested in later withdrawal [t(19)=0.814, p=0.484]; however, their responding was now significantly higher than that of VEH-infused rats tested in early withdrawal [t(17)=2.154, p=0.021]. Thus, the inhibitory effect of intra-PL NBQX infusion observed immediately following microinjection (**Figure 1A**) is transient and does not persist into the next day.

### The inhibitory effect of intra-PL NBQX is incubation-selective

To confirm that the reduction in incubated cocaine-craving observed in our initial experiment (**Figure 1A**) was specific to the cocaine-incubated state, we examined the effects of intra-PL NBQX infusion on cue-reinforced responding in rats tested on WD1. The results pertaining to the average behavior of the rats over the course of the last 3 days of the cocaine self-administration phase of the study are presented in **Table 1 (Expt. 2)**. Analyses of the number of active lever-presses [F(3,20)=0.000, p=0.992], inactive lever-presses [F(3,20)=3.689, p=0.074] and reinforcers earned [F(3,20)=0.462, p=0.507] did not indicate any significant group differences at the outset of testing.

Although no sex differences in responding were detected in our study of incubated craving (see above), females emitted more active lever-presses, overall, than males when tested in early withdrawal (**Figure 2A,B**) [Sex effects, for active lever: F(3,20)=23.681, p<0.001; for inactive lever: F(3,18)=2.358, p=0.147]. We also detected significant Treatment effects for both levers [for active lever: F(3,20)= 15.772, p=0.001; for inactive lever: F(3,18)= 5.795, p=.030], as well as significant Treatment X Sex interactions [for active lever: F(3,20)=18.071, p<0.001; F(3,18)=5.795, p=0.030]. Deconstruction of the interaction for active lever-pressing along the Sex factor indicated a robust NBQX-induced *increase* in cue-reinforced responding by female rats [t(8)=6.50, p<0.001], that was not apparent in males (**Figure 2A**) [t(8)=0.886, p=0.861], with a similar pattern of results observed for inactive lever-pressing (**Figure 2B**) [for females: t(8)=3.792, p=0.005; for males: t(6)=0.000, p=1.000]. Together with our results for incubated cocaine-craving (**Figure 1**), these data indicate that the capacity of NBQX to lower cocaine-craving is selective for the incubated state. Further, the results from this study argue that the effect of intra-PL NBQX infusion on incubated cocaine-craving does not reflect acute motor, motivational or cognitive impairing effects of AMPAR blockade within the PL.

**Figure 2:**
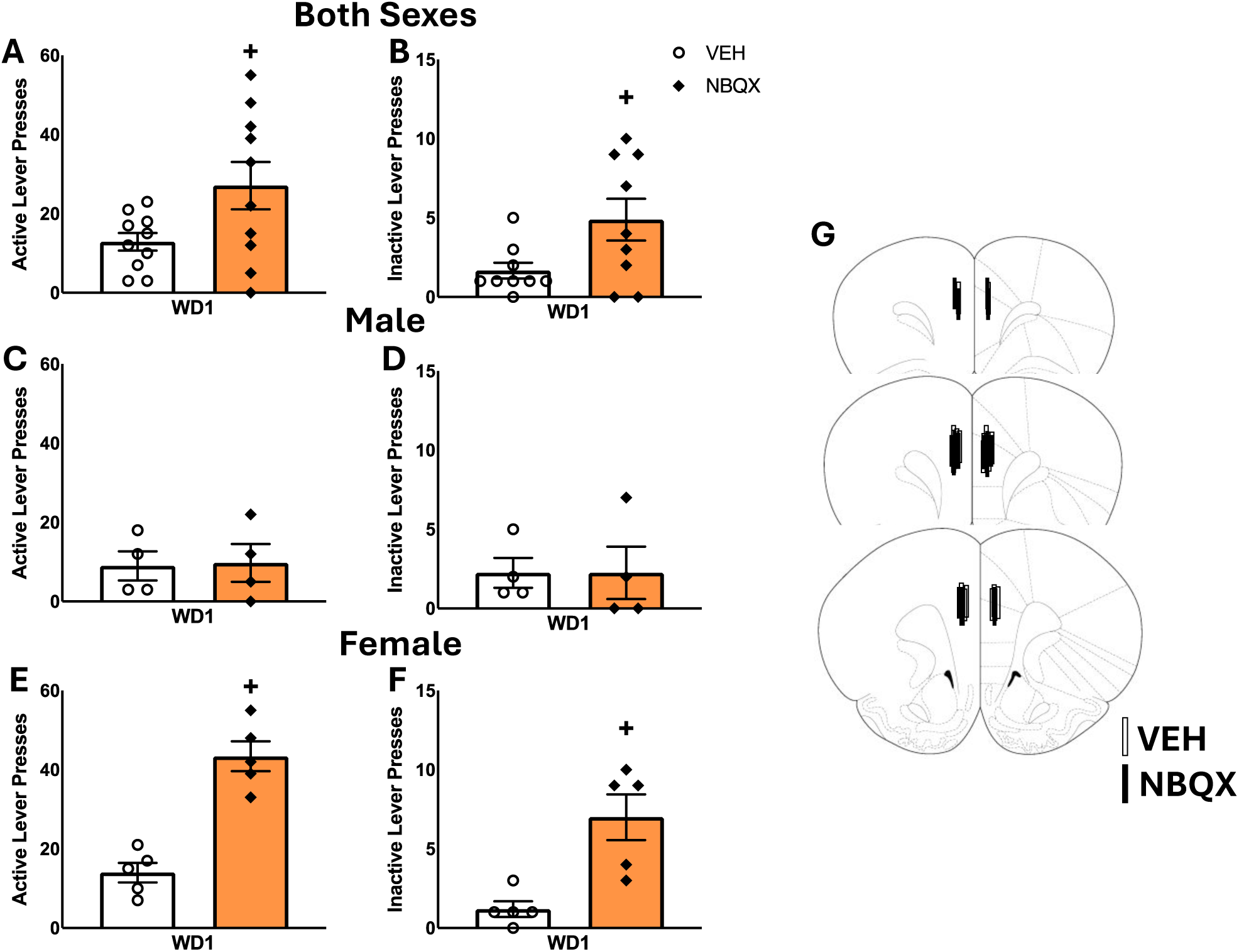
Summary of the effects of an intra-PL infusion of NBQX (1 *μ*g/side) on active and inactive lever-responding during cue tests conducted on withdrawal day 1 (WD1). As a sex difference in the effect of NBQX was detected in this study, the behavior is depicted for both sexes combined **(A,B)**, as well as for males **(C,D)** and females **(E,F)** separately. The data represent the means ± SEMs of the number of individual rats indicated. **(G)** Cartoon depicting the placements of the microinjectors within the PL. *p<0.05 vs. VEH.

### Immunoblotting for correlates of incubated cocaine-seeking

Given the effects of intra-PL NBQX infusion cue-elicited cocaine-craving and its incubation during protracted withdrawal, we next determined if AMPAR subunit expression within the PL, as well as the more ventral IL subregion, might correlate with behavior. The GluA1 subunit is the most prevalent AMPAR subunit in the brain (e.g., Rozov and Burnashev, 1999). Thus, GluA1 expression was used as a gross index of total AMPAR expression, while the GluA2 subunit was examined to assay potential changes in the number of CP-vs. CI-AMPARs (c.f., Wolf, 2025). The results pertaining to the average behavior of the male and female rats over the course of the last 3 days of the cocaine self-administration phase of the study are presented in **Table 1 (Expt. 3)**. Analyses of the number of active lever-presses [for females: F(1,27)<0.129, p’s>0.721; for males: F(1,22)<1.522, p’s>=0.230], inactive lever-presses [for females: F(1,27)<1.919, p’s>0.177; for males: F(1,22)<0.398 p’s>0.535] and reinforcers earned [for females: F(1,27)=0.000 p’s>0.985; for males: F(1,22)<2.730 p’s>0.112] did not indicate any significant group differences at the outset of testing.

#### Cocaine-seeking behavior

Significant Group X Withdrawal interactions were detected for the number of active lever-presses emitted during the 2-h cue test by both male (**Figure 3A**) and female rats (**Figure 3C**) [for males: F(1,46)=4.126, p=0.049; for females: F(1,48)=5.169, p=0.028]. For both sexes, these interactions reflected a time-dependent increase in cue-reinforced responding in the cocaine-experienced rats [for males: t(22)=3.355 p=0.0029; for females: t(26)=2.739, p=0.0110]. Cocaine-naive male controls also emitted more active lever-presses in late versus early withdrawal, but this effect was not detected in female controls [for males: t(22)=2.614,p=0.012; for females: t(18)=0.1546, p=0.879]. While no interaction was detected for inactive lever-responding by male rats (**Figure 3B**) [F(1,47)=.031, p=0.862], the interaction term was significant for females (**Figure 3D**) [F(1,48)=6.086, p=0.018] and reflected a time-dependent increase in inactive lever-responding selectively in the cocaine-experienced animals [for Controls: t(16)=0.8823, p=0.3907; for Cocaine: t(24)=2.602, p=0.016].

**Figure 3.**
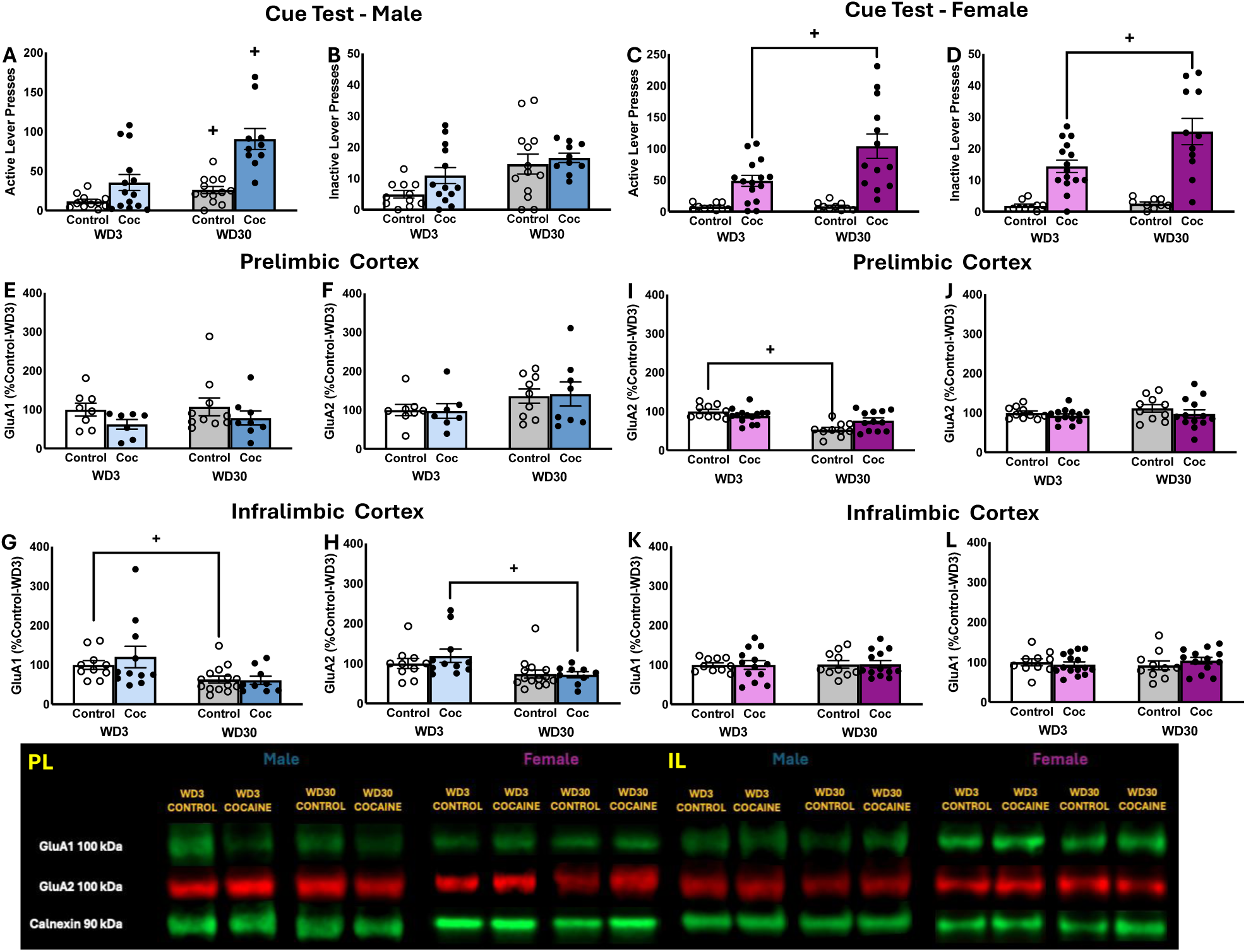
Summary of the behavior exhibited by male **(A,B)** and female **(C,D)** rats during the test for incubated cocaine-craving. Summary of the immunoblotting results for the PL of males **(E,F)** and females **(G,H)**, as well as the immunoblotting results for the IL of males **(I,J)** and females **(K,L)**. The data represent the means ± SEMs of the number of individual rats indicated. Representative immunoblots are also provided. +p<0.05 vs. WD1.

#### AMPAR subunits within mPFC subregions

A comparison of the total protein expression of GluA1 and GluA2 subunits within the PL of male rats failed to detect any differences for either subunit (**Figure 3E,F**) [for GluA1: F(3,45)<1.562, p’s>0.0218; for GluA2: (F3,45)<2.844, p’s>0.056]. A comparable analysis of AMPAR expression within the PL of female rats also failed to detect group differences in GluA1 (**Figure 3G**) [F(3,44)<1.079, p’s>0.304]. In contrast, a significant Group effect [F(3,46)=24.320, p<0.001] and Group X Withdrawal interaction [F(3,46)=.8.235, p=0.006] were detected for PL GluA2 expression in female rats (**Figure 3H**). However, deconstruction of the interaction along with Group factor indicated that this interaction reflected a time-dependent reduction in GluA2 expression in the cocaine-naive controls, with no change detected in cocaine-experienced females [for Control: t(22)=2.616, p=.016; for Cocaine: t(24)=1.428, p=0.166]. Consequently, GluA2 expression within the PL was higher in cocaine-experienced versus -naive females at the WD30 time-point.

In contrast to the PL, a time-dependent decrease in both GluA1 and GluA2 expression was observed within the IL of male rats (**Figure 3I,J**) [for GluA1: F(3,44)=8.628, p=0.005; for GluA2: F(3,44)=8.725, p=0.005]. Although it appeared that the Withdrawal effects for GluA1 and GluA2 expression were driven, respectively, by the cocaine-naive controls and the cocaine-experienced males, we detected no significant Group effects or Group X Withdrawal interactions for either subunit [for GluA1: F(3,44)<0.462, p’s>0.500; for GluA2: F(3,44)<0.696, p’s>0.409]. No changes in GluA1 or GluA2 expression were observed within the IL of female rats (**Figure 3K,L**) [for GluA1: F(3,47)<0.019, p’s>0.893; for GluA2: F(3,46)<1.694, p’s>0.199]. Taken together, these data argue that the expression of incubated cocaine-craving is not associated with changes in the total protein expression of GluA1 or GluA2 subunit expression within either mPFC subregion.

### The influence of estrous phase on incubated cocaine-craving and AMPAR expression

Although we did not detect any overt sex differences in the magnitude of incubated craving in the present study (**Figure 1**), we wanted to see if behavior and subunit expression might fluctuate with estrous phase in cocaine-experienced females as reported previously in the literature (Corbett et al., 2021; Kerstetter et al., 2008; Nicholas et al. 2019). Indeed, the number of active lever-presses varies with estrous cycle phase in cocaine-experienced females (**Figure 4A**) [α=0.1; Stage effect: F(5,28)=2.638, p=0.094; Withdrawal effect: F(5,28)=7.210, p=0.014; interaction: F(5,28)=2.918, p=0.075]. The Stage effect reflected higher responding in estrus females than those in diestrus (p=0.001) or proestrus (p=0.042), while deconstruction of the interaction along the Stage factor indicated that only estrus females exhibited higher responding on WD30 vs. WD1 [α=0.1; estrus: t(5)=2.172, p=0.082; diestrus: t(13)=1.058, p=0.309; proestrus: t(4)=0.636, p=0.560). Inactive lever-pressing behavior also varied with estrous phase (**Figure 4B**) [α=0.1;Stage effect: F(5,27)=3.532, p=0.048; Withdrawal effect: F(5,27)=6.681, p=0.017; interaction: F(5,27)=3.007, p=.071], and this Stage effect also reflected differential responding by estrus females versus diestrus (p<0.001) and proestrus females (p=0.011) and deconstruction of the interaction for inactive lever-pressing that only estrus females exhibited higher responding on WD30 vs. WD1 [estrus: t(5)=2.993, p=0.033; diestrus: t(12)=0.514, p=0.616; proestrus: t(4)=1.187, p=.301].

**Figure 4.**
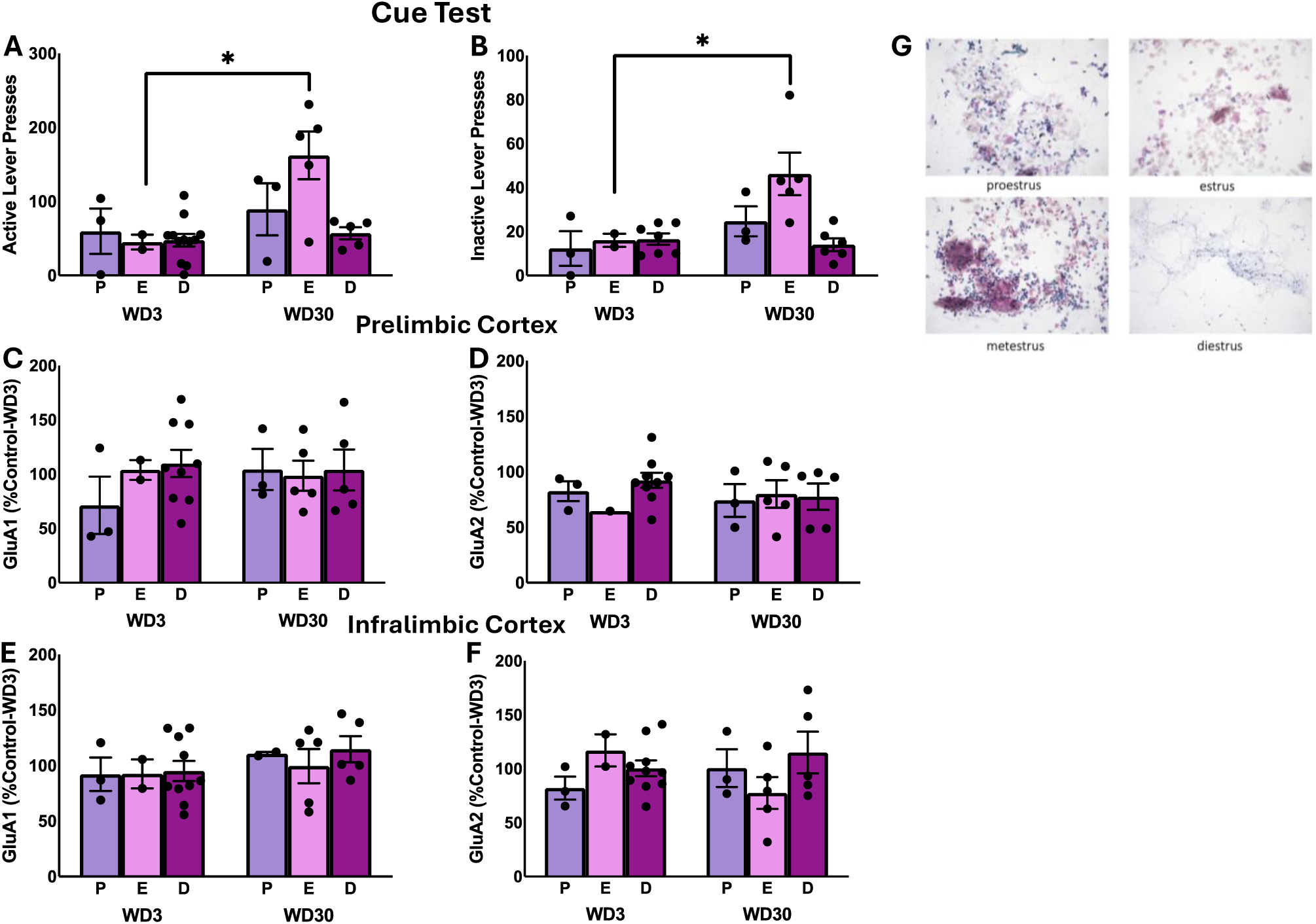
Comparison of responding of female rats in proestrus (P), diestrus (D) and estrus (E) during the test for incubated cocaine seeking **(A,B)**, as well as protein expression within the PL **(C,D)** and the IL **(E,F)**. The data represent the means ± SEMs of the number of individual rats indicated. **(G)** Representative images of the distinctions in vaginal cell cytology across the different phases of the estrous cycle.

The estrous cycle-related changes in the behavior of cocaine-experienced females were not accompanied by any overt estrous cycle-related effects on GluA1 or GluA2 expression within the PL (**Figure 4C,D**) [for GluA1: F(5,27)<1.394, p’s>0.250; for GluA2: F(5,26)<0.554, p’s>0.583] or the IL of cocaine-experienced female rats (**Figure 4E,F**) [for GluA1: F(2,27)<0.617, p’s>0.548; for GluA2: F(5,26)<1.902, p’s>0.0172].

### Immunoblotting for correlates of incubated sucrose-seeking

The results pertaining to the average behavior of the rats over the course of the last 3 days of the sucrose reinforcement phase of the study are presented in **Table 1 (Expt. 4)v** and the results of the statistical analyses of these data are described in Cano et al. (2025). The table in **Figure 5** summarizes the behavioral results from the tests for incubated sucrose-craving, in which rats of both sexes exhibited comparable incubated sucrose-craving during protracted withdrawal, as well as increased responding on the inactive lever (Cano et al., 2025). As all rats in this study were sucrose-experienced, the data are expressed relative to the male rats tested for sucrose-seeking in early withdrawal. Overall, females tended to exhibit higher GluA1 expression within the PL (**Figure 5A**) [Sex effect: F(3,47)=3.189, p=0.081; Withdrawal effect and interaction: F(3,47)<1.216, p’s>0.275] and the sex difference in GluA2 expression was statistically significant (**Figure 5B**) [Sex effect: F(3,48)=5.031, p=0.030; Withdrawal effect and interaction: F(3,47)<0.994, p’s>0.323]. When subunit expression was compared within the IL, we detected a significant Sex X Withdrawal interaction for GluA1 (**Figure 5C**) [F(3,56)=4.496, p=0.039] that reflected a time-dependent increase in GluA1 in male, but not female, sucrose-seeking rats [for males: t(26)=-2.030, p=0.053; for females: t(26)=0.930, p=0.361]. No group differences in GluA2 expression were observed within the IL (**Figure 5D**) [F(3,56)<2.018, p’s>0.161]. Thus, the expression of incubated sucrose-craving is associated with increased GluA1 expression within the IL, at least in male rats.

**Figure 5.**
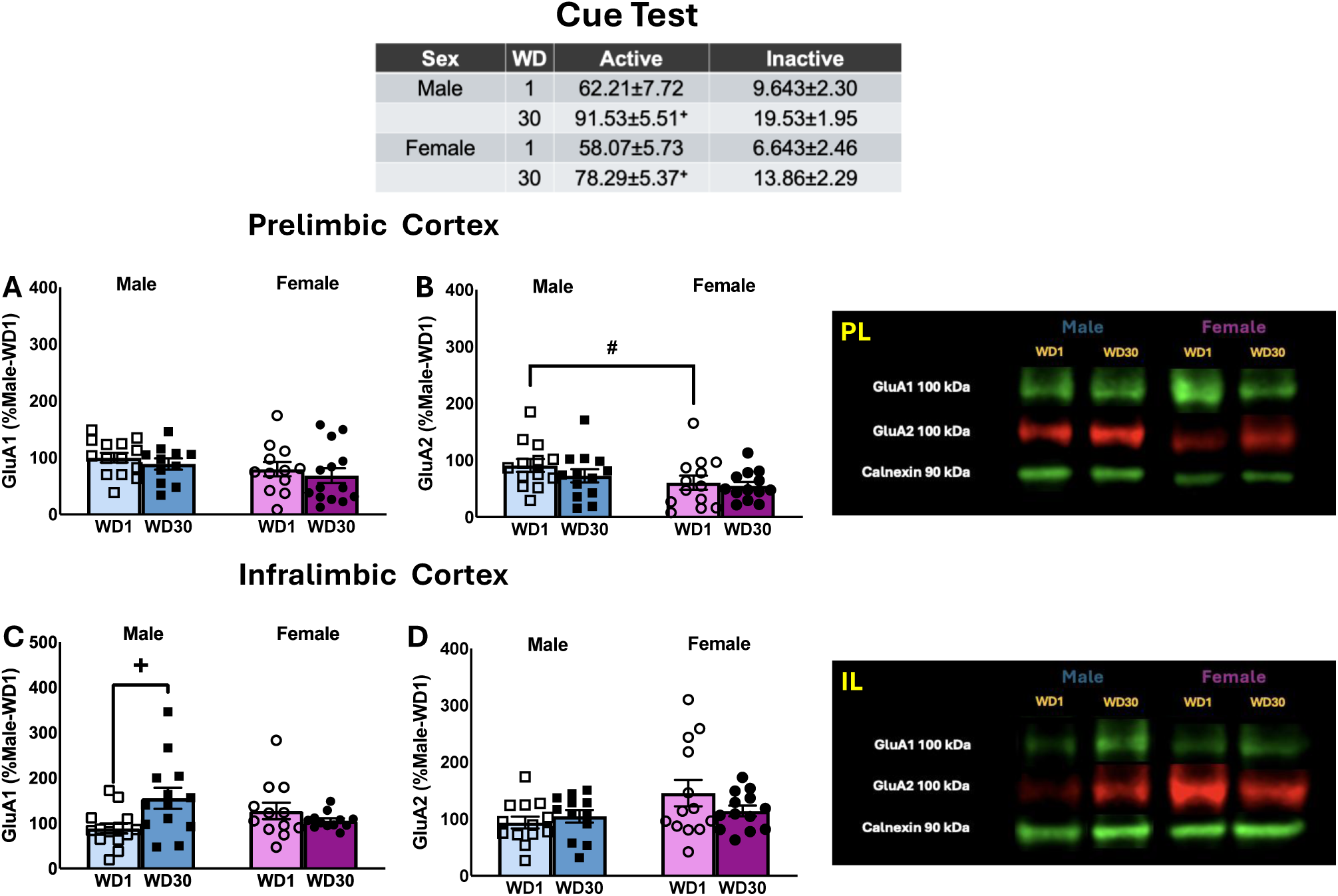
Table: Summary of the behavior exhibited by male and female rats during the test for incubated sucrose-craving, as well as the immunoblotting results for the PL **(A,B)** and IL **(C,D)**. The data represent the means ± SEMs of the number of individual rats indicated. Representative immunoblots are also provided. +p<0.05 vs. WD1; # p<0.05 male vs. female

## Discussion

AMPARs are considered critical biomolecular mediators of synaptic plasticity, including that associated with withdrawal from repeated cocaine exposure (Bowers et al., 2010; Loweth et al., 2014; Wolf, 2025). Despite this, and considerable evidence from human imaging studies implicating mPFC hyper-activation in drug cue-reactivity (e.g., Devoto et al., 2020; Goldstein & Volkow, 2011; Mohd Nawawi et al., 2024), no study to our knowledge has directly examined the role for AMPARs within the mPFC in the intensification of drug cue-reactivity that occurs during protracted cocaine withdrawal. Herein, we show that an intra-PL infusion of the AMPAR antagonist NBQX is sufficient to block the expression of incubated cocaine-craving expressed by both female and male rats tested in later withdrawal. In contrast, intra-PL NBQX infusion produces the opposite effect in female rats tested in early withdrawal and *increases* cue-reactivity. Despite these neuropharmacological results implicating AMPAR activation within the PL as important for modulating cue-elicited cocaine-craving, we failed to detect any cocaine- or cocaine incubation-related changes in GluA1 or GluA2 subunit expression within either the PL or IL subregion under conventional immunoblotting procedures using whole-cell lysates. In contrast, the expression of incubated sucrose-craving was associated with an increase in GluA1 expression within the IL of male rats only. Below, we discuss these findings within the context of the limited literature focused on the role for glutamate transmission within the mPFC in cocaine- and sucrose-craving, as well as their incubation during protracted reinforcer abstinence.

### Inhibition of PL AMPARs blocks incubated cocaine-craving and inhibitory effect is not present in early withdrawal

Incubated cocaine-craving is associated with a cue-elicited increase in extracellular glutamate within the mPFC (Shin et al., 2016) and this glutamate release, particularly within the PL subregion, is necessary for incubated cocaine-craving (Shin et al., 2018). The current studies demonstrate that AMPAR inhibition within the PL by NBQX blocks incubated cocaine-craving (**Figure 1**). Our collective findings argue that cocaine cue-elicited glutamate release during late withdrawal activates AMPARs within the PL to drive incubated cocaine-craving. Thus, as reported for the NAc (Conrad et al., 2008; Cornish and Kalivas 2000; Loweth et al., 2014), AMPAR stimulation within the mPFC also plays a necessary role in the expression of cocaine-craving following a period of cocaine-abstinence.

Herein, the effect of AMPAR inhibition within the PL on cocaine-seeking was selective for the incubated state as NBQX infusion did not reduce responding in early withdrawal (**Figure 2**). Quite opposite, AMPAR inhibition increased cue-elicited seeking selectively in female rats tested on WD1 (**Figure 2C-F**), indicating that AMPAR activation during early withdrawal normally serves to blunt the motivational properties of cocaine-related cues. As this study employed a single, relatively high, dose of NBQX (see Russell et al., 2016), it remains to be determined whether the female-selective effect of NBQX on WD1 reflects sex differences in the affinity of AMPARs for NBQX or baseline activity of mPFC AMPARs. Nevertheless, our neuropharmacological studies using the 1.0 *μ*g/side NBQX dose argue that AMPARs within mPFC undergo some form of plasticity over the course of cocaine withdrawal that changes the functional import of receptor activation for cocaine-seeking behavior. While not yet assayed within a model of incubated cocaine-craving, a time-dependent insertion of CP-AMPARs are reported to occur within layer 5 of mPFC pyramidal neurons of cocaine-sensitized mice during withdrawal. This CP-AMPAR insertion impairs normal mGlu1/mTOR-dependent long-term depression in this region and induces a “malplastic state” (Ruan and Yao, 2021). Indeed, prior neuropharmacological studies from our group have implicated both PI3K/Akt/mTOR activation (Chiu et al., 2021; Szumlinski et al., 2019) and lowered mGlu1 function within mPFC (Ben-Shahar et al., 2013) as critical molecular adaptations for the expression and persistence of incubated cocaine-craving, respectively. Now that we have confirmed a selective role for AMPAR stimulation within the PL for incubated cocaine-seeking, future studies will seek to determine the contribution of CP-AMPARs by testing the effects of the selective GluA2-lacking AMPAR antagonist Naspm. As inhibition of glutamate release within the IL also dampens incubated cocaine-craving (Shin et al., 2018), it will be important also to assay the relative role for different AMPAR subtypes expressed within the IL as driving cocaine-craving in the short and longer term. Finally, the inhibitory effect of intra-NAc Naspm extends from models of incubated cocaine-craving (e.g., Conrad et al., 2008; Kawa et al., 2022; Loweth et al., 2014), to those of methamphetamine- and oxycodone-craving (Nicholas et al., 2019; Scheyer et al., 2016; Wong et al., 2023), raising the possibility that withdrawal-dependent insertion of CP-AMPARs may also contribute to perturbations in synaptic plasticity within mPFC produced by other drugs of abuse.

### No overt changes in GluA1 and GluA2 expression associated with incubated cocaine-craving

Prior immunoblotting studies by our group indicated that the detection and magnitude of incubation-related changes in the expression of certain proteins within mPFC (e.g., Akt activation, mGlu1, mGlu5 and Homer2) are more robust in rats with a prior history of extended-versus shorter-access to intravenous cocaine (e.g., Chiu et al., 2021; Huerta Sanchez et al., 2023 vs. Ben-Shahar et al., 2013; Gould et al., 2015; Szumlinski et al., 2019). Indeed, evidence also indicates that the duration of daily cocaine-access impacts the ability to detect CP-AMPAR-related changes, at least within the NAc (Purgianto et al., 2013). However, the null results for GluA1 and GluA2 expression within the mPFC of the cocaine-incubated rats in the present study (**Figure 3**) align with those reported previously for rats expressing incubated cocaine-craving following shorter-access self-administration paradigms (Huerta Sanchez et al., 2023), arguing against the duration of cocaine-access as being a critical factor in our inability to detect changes in AMPAR subunit expression.

Prior studies of AMPAR subunit expression within the NAc have reported increased cell surface *and* intracellular GluA1 expression in cocaine-incubated rats (Conrad et al., 2008) that is purported to reflect up-regulated GluA1 translation (e.g., Hwang et al., 2025; Scheyer et al., 2014; Stefanik et al., 2018). Based on the results of Conrad et al. (2008), we rationalized that if incubated cocaine-craving was associated with increased surface and intracellular GluA1 expression, then our conventional immunoblotting procedures, conducted on whole-cell lysates, would be sufficient to detect changes in subunit expression within mPFC if they occurred. Indeed, we successfully detected a time-dependent increase in GluA1 expression within the IL of male rats exhibiting incubated sucrose-seeking (**Figure 5C**). Thus, it may be that (1) incubated cocaine-craving is completely dissociated from changes in AMPAR translation/expression within mPFC or (2) the changes in AMPAR expression associated with the cocaine-incubated state are too subtle to detect in whole-cell lysates. In light of electrophysiological evidence supporting the insertion of GluA2-lacking AMPARs within mPFC pyramidal neurons during cocaine withdrawal (Run and Yao, 2021), we propose that any future immunoblotting studies of AMPAR subunit expression within mPFC employ subcellular fractionation or biotinylation procedures to isolate cell surface subunit expression to better inform the relationship between incubated cocaine-craving and the subunit composition of functionally relevant AMPARs.

### Estrous phase influences cue-induced cocaine-craving but not AMPAR expression

The magnitude of incubated cocaine-craving by female rats can be influenced by their hormonal status with heightened craving observed in the estrus phase compared to both females in non-estrus phases and males (Corbett et al., 2021; Kerstetter et al., 2013; Nicolas et al., 2019). Consistent with these prior studies, the female rats identified in the present study as being in estrus exhibited more cue-induced cocaine-seeking, compared to females rats in diestrus or proestrus (**Figure 4A-B**). How the estrous cycle impacts the biomolecular correlates of incubated cocaine-craving is not known. As such, we examined how GluA1 and GluA2 expression might vary with cycle phase but failed to detect differences in subunit expression as a function of estrous cycle phase, at least in cocaine-experienced rats (**Figure 4C-F**). Whether or not this failure to detect an estrous cycle effect on AMPAR expression reflects our examination of whole-cell lysates or a genuine lack of any relationship in cocaine-experienced females cannot be discerned from the design of the present study as vaginal cytology was not conducted on our cocaine-naive control females.

### Sex-selective protein correlates of incubated sucrose-seeking

The similar temporal profile of incubated craving for different drugs of abuse and non-drug reinforcers (e.g., sucrose, saccharin and high-fat foods) have led to the theory that common neurocircuitry and biomolecular changes might underpin the phenomenon of incubated craving (c.f., Grimm, 2020). Indeed, several studies have examined for common biomolecular mechanisms in the incubation of craving for drug versus non-drug reinforcers (Knackstedt & Kalivas, 2014; Blanco-Gandia et al., 2020; Venniro et al., 2021). Of relevance to the present study, increased indices of neuronal activity are reported within the PL and IL of cocaine-, heroin-, and sucrose-experienced rats (Koya et al, 2009; Counotte et al., 2013; Grimm et al., 2016), suggesting that these mPFC subregions may be components of a common neurocircuitry driving incubated cue-elicited reward-seeking. However, the neurochemical data to date indicate that the capacity of sucrose-associated cues to elevate extracellular glutamate levels within the mPFC dissipates, rather than intensifies, in male rats with the passage of time in withdrawal (Shin et al., 2016). Thus cocaine- and sucrose-craving during protracted abstinence are associated with opposite time-dependent changes in mPFC extracellular glutamate, which would be predicted to induce distinct changes in glutamate-related signaling in mPFC subregions. Indeed, a recent study by our group failed to identify “cocaine-like” changes in several glutamate-related proteins within mPFC subregions of rats expressing incubated sucrose-craving (Cano et al., 2025 vs. Ben-Shahar et al., 2013; Chiu et al., 2021; Gould et al., 2015; Miller et al., 2016; Szumlinski et al., 2019). Aligning with these discrepancies in protein expression, we detected no changes in PL or IL levels of either AMPAR subunit in cocaine-incubated rats, while a time-dependent increase in GluA1 expression was detected within the IL of male, but not female, rats (**Figure 5C**).

As our prior microdialysis study employed males only (Shin et al., 2016), we do not know whether females also exhibit a time-dependent reduction in sucrose cue-reactivity. However, it is worth noting that the male-selectivity of the observed change in GluA1 expression within the IL of sucrose-incubated males align with our recent finding that incubated sucrose-craving *in these same males* is associated with increased IL expression of p(Ser473)-Akt, p(Ser729)-PKCɛ, and p(Ser2448)-mTOR (Cano et al., 2025), arguing that the present result for IL GluA1 expression is not likely spurious. In contrast to males, the same female rats as those employed in the present study exhibit lower total protein expression of p(Ser473)-Akt and p(Ser729)-PKCɛ within the PL (Cano et al., 2025). Thus, even though the magnitude of incubated sucrose-craving is comparable between male and female rats (**Figure 5**; Cano et al., 2025), our immunoblotting data to date indicate that the biomolecular mechanisms (at least within mPFC) associated with incubated sucrose-craving are not only sex-dependent, but distinct from those observed in animals expressing incubated cocaine-craving. As an increase in GluA1 subunit expression is associated with the insertion of CP-AMPARs in NAc (e.g., Conrad et al., 2008), an important goal for future work is to determine whether sex differences exist in the effects of the GluA2-lacking AMPAR antagonist Naspm on the expression of incubated sucrose-craving.

### Conclusions

AMPAR inhibition within the PL blocked incubated cocaine-craving during protracted withdrawal in rats of both sexes, while increasing cocaine cue-reactivity in female rats during early withdrawal. Although AMPAR activation within the PL is clearly necessary for the expression of incubated cocaine-craving, incubated cocaine-craving is not overtly related to GluA1 or GluA2 subunit expression within either the PL or IL. In contrast, increased GluA1 expression within the IL is associated with incubated sucrose-craving, but in male rats only. These data indicate a key role for PL AMPARs in driving incubated cocaine-craving and suggest that AMPARs within the IL may potentially gate the development of incubated sucrose-craving in males. Our findings inform as to the biomolecular mechanisms within mPFC that drive incubated craving across drug and non-drug reinforcers of relevance to both the efficacy and side-effect profiles of glutamate-targeting therapies for treating pathological craving in both cocaine use and eating disorders.

## Data Availability Statement

The raw data supporting the conclusions of this article will be made available by the authors, without undue reservation.

## Ethics Statement

The animal study was reviewed and approved by Institutional Animal Care and Use Committee of the University of California Santa Barbara.

## Author Contributions

The authors contributed to this report in the following ways. Conceptualization, L.L.H.S., M.G.T, H.H.T.D , and K.K.S.; formal analysis, K.K.S., and L.L.H.S..; investigation, L.L.H.S., M.G.T., H.H.T.D., S.V.V., S.C.R., T.L.L., P.B.J., A.Y.N., and F.J.C.; writing—original draft preparation, L.L.H.S. and K.K.S.; writing—review and editing, L.L.H.S., M.G.T., H.H.T.D., S.V.V, S.C.R., T.L.L., P.B.J., A.Y.N., F.J.C., T.E.K., and K.K.S; visualization, L.L.H.S. and K.K.S.; supervision, T.E.K. and K.K.S.; project administration, L.L.H.S. and K.K.S; funding acquisition, L.L.H.S., M.G.T., S.C.R., T.L.L., K.K.S and T.E.K. All authors have read and agreed to the published version of the manuscript.

